# BPG: Seamless, Automated and Interactive Visualization of Scientific Data

**DOI:** 10.1101/156067

**Authors:** Christine P’ng, Jeffrey Green, Lauren C. Chong, Daryl Waggott, Stephenie D. Prokopec, Mehrdad Shamsi, Francis Nguyen, Denise Y.F. Mak, Felix Lam, Marco A. Albuquerque, Ying Wu, Esther H. Jung, Maud H.W. Starmans, Michelle A. Chan-Seng-Yue, Cindy Q. Yao, Bianca Liang, Emilie Lalonde, Syed Haider, Nicole A. Simone, Dorota Sendorek, Kenneth C. Chu, Nathalie C. Moon, Natalie S. Fox, Michal R. Grzadkowski, Nicholas J. Harding, Clement Fung, Amanda R. Murdoch, Kathleen E. Houlahan, Jianxin Wang, David R. Garcia, Richard de Borja, Ren X. Sun, Xihui Lin, Gregory M. Chen, Aileen Lu, Yu-Jia Shiah, Amin Zia, Ryan Kearns, Paul C. Boutros

**Affiliations:** Informatics & Biocomputing Program, Ontario Institute for Cancer Research, Toronto, Canada; Department of Medical Biophysics, University of Toronto, Toronto, Canada; Department of Pharmacology and Toxicology, University of Toronto, Toronto, Canada

**Author notes:** Present address: Center for Computational Research, Buffalo Institute for Genomics and Data Analytics, NYS Center for Excellence in Bioinformatics & Life Science, University at Buffalo. Email addresses: CP, LCC, SDP, FN, FL, YW, MHWS, CQY, EL, NAS, KCC, NSF, NJH, ARM, JW, RdB, XL, AL, AZ, JG, DW, MS, DYFM, MAA, EHJ, MAC, BL, SH, DS, NM, MG, CF, KEH, DRG, RXS, GMC, YS RK. Corresponding author contact information: Paul C. Boutros; Ontario Institute for Cancer Research, MaRS Centre, 661 University Avenue, Suite 510, Toronto, Ontario, Canada, M5G 0A3.

## Abstract

We introduce BPG, an easy-to-use framework for generating publication-quality, highly-customizable plots in the R statistical environment. This open-source package includes novel methods of displaying high-dimensional datasets and facilitates generation of complex multi-panel figures, making it ideal for complex datasets. A web-based interactive tool allows online figure customization, from which R code can be downloaded for seamless integration with computational pipelines. BPG is available at http://labs.oicr.on.ca/boutros-lab/software/bpg

---

Biological experiments are increasingly generating large, multifaceted datasets. Exploring such data and communicating observations is, in turn, growing more difficult and the need for robust scientific data-visualization is growing rapidly^1,2,3,4^. Myriad data visualization tools exist, particularly as web-based interfaces and non-R-based local software packages. Unfortunately these do not integrate easily into R-based statistical pipelines such as the widely used Bioconductor^5^. Within R, visualization packages exist, including base graphics^6^, ggplot2^7^, lattice^8^, Sushi^9^, circlize^10^, multiDimBio^11^, NetBioV^12^, GenomeGraphs^13^ and ggbio^14^. These lack publication-quality defaults, contain limited plot types, provide limited scope for automatic generation of multi-panel figures, are constrained to specific data-types and do not allow interactive visualization.

Good visualization software must create a wide variety of chart-types in order to match the diversity of data-types available. It should provide flexible parametrization for highly customized figures and allow for multiple output formats while employing reasonable, publication-appropriate default settings, such as producing high resolution output. In addition, it should integrate seamlessly with existing computational pipelines while also providing an easily intuitive, interactive mode. There should be an ability to transition between pipeline and interactive mode, allowing cyclical development. Finally, good design principles should be encouraged, such as suggesting appropriate color choices and layouts for specific use-cases. To help users quickly gain proficiency, detailed examples, tutorials and an application programming interface (API) are required. To date, no existing visualization suite fills these needs.

To address this gap, we have created the BPG library, which is implemented in R using the grid graphics system and lattice framework. It generates a broad suite of chart-types, ranging from common plots such as bar charts and box plots to more specialized plots, such as Manhattan plots (**Figure 1;** code is in **Supplementary File 1**). These include some novel plot-types, including the dotmap: a grid of circles inset inside a matrix, allowing representation of four-dimensional data (**Figure 1n**). Each plotting function is highly parameterized, allowing precise control over plot aesthetics. The default parameters for BPG produce high resolution (1600 dpi) TIFF files, appropriate for publication. The file type is specified simply by specifying a file extension. Other default values contribute to graphical consistency including: the inclusion of tick marks, selection of fonts and suggested default colors that work together to create a consistent plotting style across a project.

**Figure 1.**
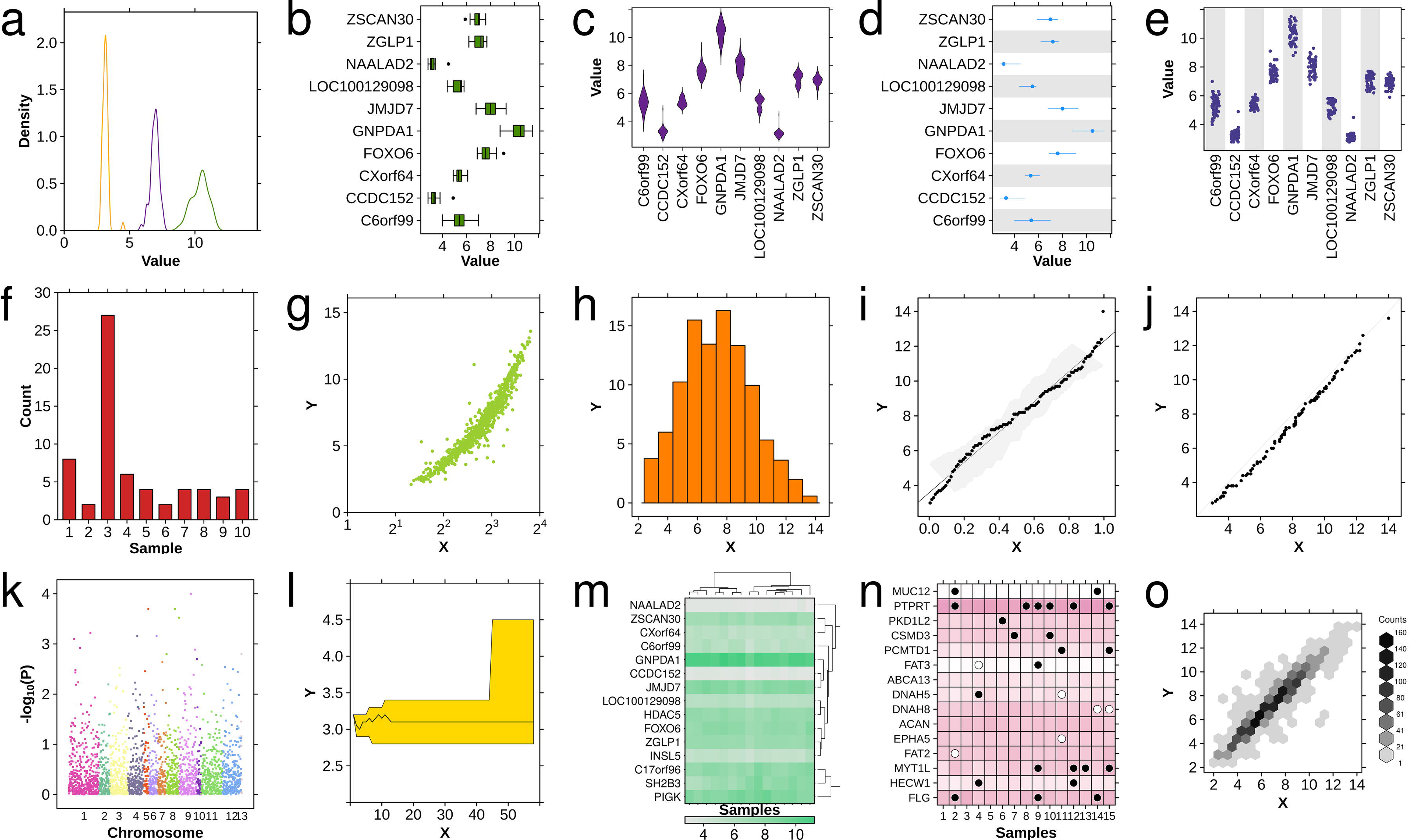
Available chart-types. The basic chart-types available in BPG: (a) density plot, (b) boxplot, (c) violin plot, (d) segplot, (e) strip plot, (f) barplot, (g) scatterplot, (h) histogram, (i) qqplot fit, (j) qqplot comparison, (k) Manhattan plot, (l) polygon plot, (m) heatmap, (n) dotmap and (o) hexbinplot. All plots are based upon the datasets included in the BPG package and code is given in **Supplementary File 1**.

Default values have been optimized to generate high-quality figures, reducing the need for manual tuning. However, even good defaults will not be appropriate for every use-case^15^. **Supplementary Figure 1** demonstrates a single scatter plot created using four separate graphics frameworks with either default or optimized settings: BPG, base R graphics, ggplot2, and lattice. BPG required half as much code as the other frameworks for both default and optimized plots, with equal or higher quality (**Supplementary File 2**).

To facilitate rapid graphical prototyping, an online interactive plotting interface was created (http://bpg.oicr.on.ca). This interface allows users to easily and rapidly see the results of adjusting parameter values, thereby encouraging precise improvement of plot aesthetics. The R code generated by this interface is also made available for download, as can a methods paragraph allowing careful reporting of plotting options. A public web-interface is available, and local interfaces can be easily created.

One critical feature of BPG is the ability to combine multiple plots into a single figure: a technique used widely in publications. This is accomplished by the `create.multiplot` function, which automatically aligns plots and standardizes parameters such as line widths and font sizes within the final figure. This replaces the slow and error-prone manual combination of figures using PowerPoint, LaTeX or other similar software. The necessity of combining multiple plots arises from the complexity of datasets – with high dimensional data, it is often difficult to convey all relevant information within a single chart-type. Combining multiple chart-types allows more in depth visualization of the data. For example, one plot might convey the number of mutations present in different samples; a second plot could add the proportion of different mutation types, while a third could give sample-level information (**Figure 2**).

**Figure 2.**
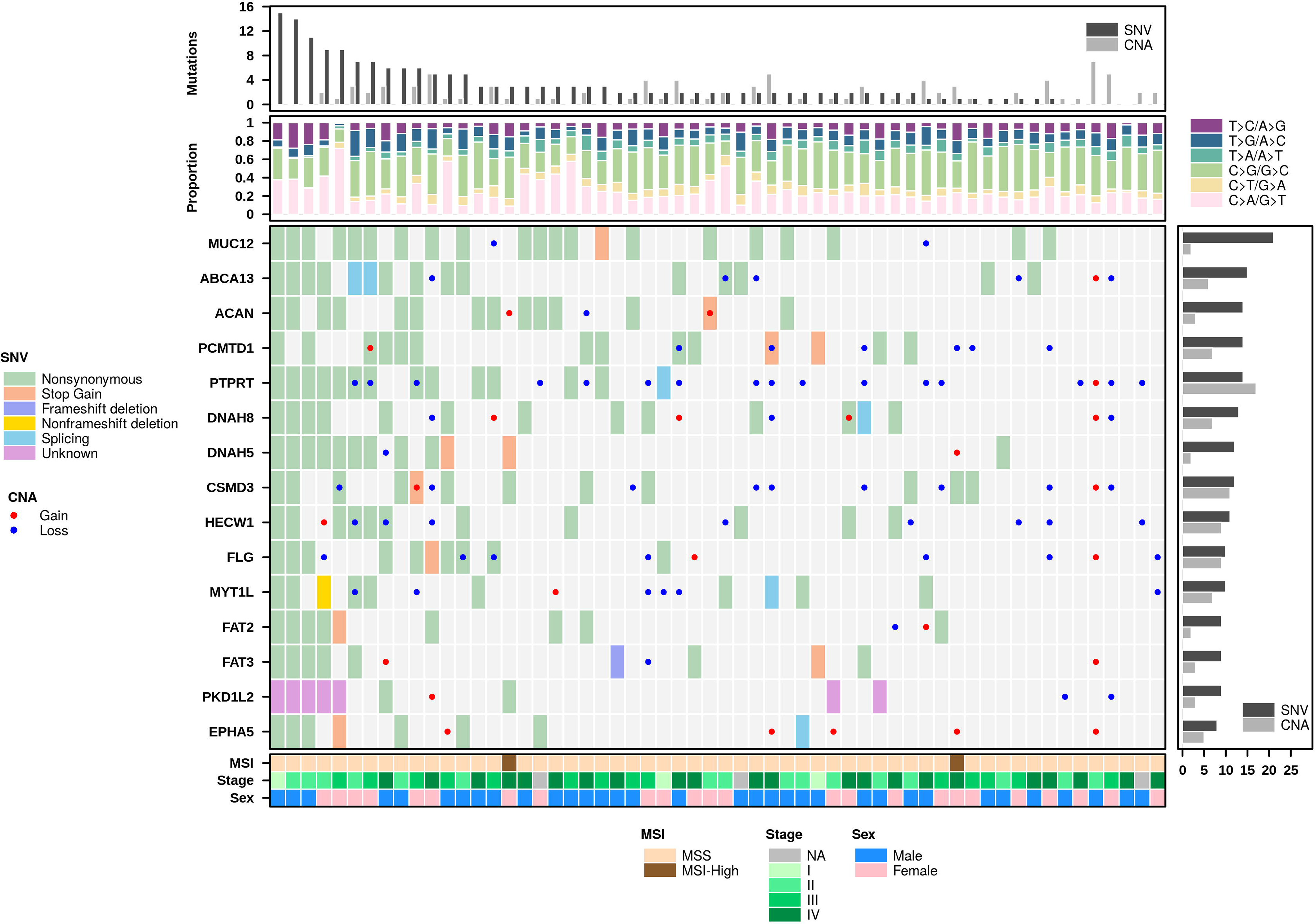
Multiplot example. The `create. multiplot` function is able to join multiple chart-types together into a single figure. In this example, a dotmap conveys the mutations present in a selection of genes (y-axis) for a number of samples (x-axis), while adjacent barplots and heatmaps provide additional information. Code used to generate this figure is available in **Supplementary File 3**.

A number of utility functions in BPG assist in plot optimization, such as producing legends and covariate bars, or formatting text with scientific notation for p-values. One difficult step in creating figures is the selection of color schemes that are both pleasing and interpretable^16,17^. BPG provides a suite of 45 color palettes including qualitative, sequential, and diverging color schemes^18^, shown in **Supplementary Figure 2**. Many optimized color schemes exist for numerous use cases including tissue types, chromosomes and mutation types. The default.colors function produces a warning when a requested color scheme is not grey-scale compatible, a common concern for figures reproduced in black and white. This is determined by converting each color to a grey value between 1 and 100, and indicating differences of <10 as not grey-scale compatible to approximate a color scheme’s visibility when printed in grey-scale. To facilitate reproducibility, image metadata is automatically generated for all plots, creating descriptors such as software and operating system versions.

Extensive documentation is provided to help new users learn how to use BPG. In order to assist researchers in determining which chart-type is appropriate for their dataset, we provide plotting examples in the documentation which are derived from a real dataset and a plotting guide is included to explain the intended use-case of each function. This guide also contains explanations of typography, basic color theory and layout design which help to improve the design of figures^19^. In addition, an online API is available with both simple and complex use-case examples for each plot-type to help users quickly learn the range of functionality available.

BPG has been used in over 60 publications to date (**Supplementary Table 1**). These plotting functions have been integrated into numerous R analysis pipelines for automated figure generation as part of the analysis of large–omic data. The plots created by this package are reproducible and maintain a consistent aesthetic. We expect that BPG will greatly facilitate improved visualization and communication of complex datasets.

## Acknowledgements

This study was conducted with the support of the Ontario Institute for Cancer Research to PCB through funding provided by the Government of Ontario. This work was supported by Prostate Cancer Canada and is proudly funded by the Movember Foundation - Grant number RS2014-01. PCB was supported by a Terry Fox Research Institute New Investigator Award and a Canadian Institutes of Health Research (CIHR) New Investigator Award. This research is funded by the Canadian Cancer Society (grant number 702528). This work was supported by the Discovery Frontiers: Advancing Big Data Science in Genomics Research program, which is jointly funded by the Natural Sciences and Engineering Research Council (NSERC) of Canada, the Canadian Institutes of Health Research (CIHR), Genome Canada, and the Canada Foundation for Innovation (CFI). This project was supported by Genome Canada through a Large-Scale Applied Project contract to PCB and Drs. Sohrab Shah and Ryan Morin. This study was conducted with the support of the Ontario Genomics Institute (to CP), the Canadian Breast Cancer Foundation (to CQY), the Ontario Graduate Scholarship (to EL), the Oncology Research and Methods Training Program (to YW, NCM and XL), the Canadian Institutes of Health Research (CIHR) (to EL, KEH, NSF and GMC), the Medical Biophysics Excellence University of Toronto Fund Scholarship (to NSF) and the CTMM framework (AIRFORCE project) and EU 7^th^ framework program (ARTFORCE) to MHWS. The authors thank all members of the Boutros lab for their assistance in testing and improving the software and our many collaborators for their suggestions and support, particular Drs. Robert Bristow, Michael Fraser, Raimo Pohjanvirta, Allan Okey and Linda Penn. We thank the OICR webdev and systems teams for support, particularly Joseph Yamada and Rob Naccarato.

### Disclosure Declaration

All authors declare that they have no competing interests.

### Author contributions

PCB conceived of the project. All authors wrote software, documentation and debugged. FM and NAS developed the interactive plotting method. CP wrote the manuscript, which all authors edited and approved.

---

**Supplementary Figure 1 | Comparison of graphical software options in R**

(a-b) are created with base R graphics, (c-d) are created using ggplot2, (e-f) are made in lattice and (g-h) use BPG. The first plot in each pair uses default settings, while the second plot has been adjusted for font sizes, axes ranges, tick mark locations, grid lines, diagonal lines, background shading and highlighted datapoints. The number of lines of code used to create default plots are: 10 for base R, 10 for ggplot2, 14 for lattice, and 5 for BPG. The customized plots use 73 lines for base R, 83 for ggplot2, 86 for lattice, and 42 for BPG. Code for generating this figure is provided in **Supplementary File 2**.

**Supplementary Figure 2 | Color palettes**

Color palettes are provided using the `default .colors` function for (a) generic use-cases and `force.color.scheme` for (b) specific use-cases. This display is generated using the `show.available.palettes` function. Interactive display of colors is also available using the `display.colors` function.

**Supplementary Table 1 | Publications using BPG**

**Supplementary File 1 | Code to generate figure 1**

**Supplementary File 2 | Code to generate supplementary figure 1**

**Supplementary File 3 | Code to generate figure 2**

